# Phage therapy minimally affects the water microbiota in an Atlantic salmon (*Salmo salar*) rearing system while still preventing infection

**DOI:** 10.1101/2023.07.03.547070

**Authors:** Alexander W. Fiedler, Madeleine S. Gundersen, Toan P. Vo, Eivind Almaas, Olav Vadstein, Ingrid Bakke

**Affiliations:** Norwegian University of Science and Technology, Trondheim, Norway

## Abstract

Excessive usage of antibiotics threatens the bacterial diversity in the microbiota of animals. An alternative to antibiotics that has been suggested to not disturb the microbiota is (bacterio)phage therapy. In this study, we challenged germ-free and microbially colonized yolk sac fry of Atlantic salmon with *Flavobacterium columnare* and observed that the mere presence of a microbiota protected the fish against lethal infection. We then investigated the effect of phage-or oxytetracycline treatment on fish survival and rearing water bacterial community characteristics using 16S rRNA gene amplicon sequencing. Phage treatment led to an increased survival of *F. columnare-*challenged fish and reduced the relative amounts of the pathogen in the water microbiota. In the absence of *F. columnare*, phage treatment did not affect the composition or the α-diversity of the rearing water microbiota. In the presence of the phage’s host, phage treatment induced minor changes to the bacterial community composition, without affecting the α-diversity. Surprisingly, oxytetracycline treatment had no observable effect on the water microbiota and did not reduce the relative abundance of *F. columnare* in the water. In conclusion, we showed that phage treatment prevents mortality while not negatively affecting the rearing water microbiota, thus suggesting that phage treatment may be a suitable alternative to antibiotics. We also demonstrated a protective effect of the microbiota in Atlantic salmon yolk sac fry.

## INTRODUCTION

Animal hosts benefit greatly from microbial colonization which provides nutrients for growth, supports normal development^[1,2]^ and protects against infections^[3–7]^. These beneficial functions are threatened by a decline in microbial diversity, leaving the organism more vulnerable to infection^[8,9]^. One major reason for this decline is the use of antibiotics^[10]^. Antibiotics disturb the microbiota, leading to decreased protection against new infections or to immediate secondary infections^[11]^. Furthermore, the overuse of antibiotics has resulted in the selection of antibiotic-resistant strains, thereby diminishing the efficacy of these drugs^[12]^.

One of the most used antibiotics for animal production in Europe is oxytetracycline (OTC)^[13]^. This broad-spectrum antibiotic binds to the 30S ribosomal subunit which inhibits bacterial protein synthesis ^[14,15]^. Due to its widespread industrial use in aquaculture, OTC has been found in ecosystems close to fish farms, such as sediments, water bodies and in aquatic organisms^[16–18]^. Already low concentrations of OTC can negatively affect aquatic organisms and their microbiota^[19–22]^ and can lead to an increase in antibiotic resistance ^[23]^. Moreover, similar to other antibiotics, OTC disturbs microbial communities, resulting in long-term changes in the community characteristics^[24]^.

Bacteriophage therapy, which employs bacteriophages to eliminate bacterial pathogens, is an encouraging substitute for antibiotics ^[25]^. Bacteriophages (phages) represent a distinct group of viruses that selectively infect and lyse bacterial cells. Given their inherent specificity, virulent phages are promising therapeutic agents to combat bacterial diseases^[26]^. Currently, phage therapy is not commonly used to treat human infections on a global level, however it is increasingly used in aquaculture ^[27]^. Due to their high specificity down to the bacterial strain level, phages are assumed to minimally influence bacterial communities. Some studies confirm this assumption (e.g.^[28–30]^), however, others have found that phage therapy affects the host-associated microbiota, likely through secondary effects due to lysis of the host bacterium^[31–33]^.

A potential target for phage therapy is columnaris disease, which is a major concern in aquaculture, especially for warm-water salmonid species ^[34,35]^. This disease is caused by the bacterium *Flavobacterium columnare*, which infects, amongst others, salmonid fish ^[4,36,37]^. While antibiotics is the common line of treatment, bacteriophage therapy against *F. columnare* in aquaculture systems has been demonstrated with promising results^[38–40]^. Here, we used a germ-free model of Atlantic salmon yolk sac fry to investigate the protective effect of the commensal microbiota against infection of *F. columnare*. We therefore expected to find higher mortality in germ-free fish challenged with the pathogen in comparison to microbially colonized fish that were challenged. We further used phage therapy against *F. columnare* and investigated its effect on the water bacterial community and on the survival of the fish in comparison to the antibiotic OTC. Here, we expected that phage treatment would have minimal impact on the bacterial communities, especially when the pathogen is absent. Contrastingly, we expected that antibiotic treatment changes the bacterial community structure and reduces α-diversity, in contrast to phage therapy.

## MATERIAL AND METHODS

### Designs of challenge experiments

Three experiments were conducted between fall 2021 and summer 2022 where fish were challenged with a bacterial pathogen. The first experiment was performed with the purpose of determining the optimal temperature for infecting Atlantic salmon yolk sac fry (*Salmo salar*) with the bacterial pathogen *Flavobacterium columnare strain* Fc7. All fish used in Exp.1 were raised germ-free (see below). Groups of 15 fish were reared in 250 ml cell culture flasks with aerated caps (hereafter referred to as “fish flasks”) at 5.6 ± 0.4 °C until 5 weeks post hatching (wph; hatching day is when 70% of all eggs have hatched). At 5 wph, the temperature in the fish flasks was gradually increased to either 10 or 14 °C over the course of one week or were kept at 6 °C (five replicate flasks per temperature and 15 flasks in total). At 6 wph and for each temperature group, three flasks were challenged with *F. columnare* Fc7, one was exposed to the none-infectious fish commensal *Janthinobacterium* sp. 3.108 and one was kept as a non-challenged control. The mortality in each flask was checked at least two times daily until the experiment was terminated at 10 days post challenge (dpc).

In experiment 2 (Exp.2), we challenged Atlantic salmon yolk sac fry with *F. columnare* Fc7 at 10 °C and subsequently added bacteriophages or oxytetracycline to the water daily. We aimed to evaluate the impact of phage-and antibiotic treatment on fish survival and the bacterial community of both the rearing water and the fish. By comparing the survival after challenge with *F. columnare* Fc7 between germ-free and microbially colonized fish (i.e., fish that were hatched under germ-free conditions but were then re-colonized by bacteria; see below), we investigated whether the microbiota protected the yolk sac fry. Both germ-free and colonized fish were raised until 5 wph at 5.2 ± 0.4 °C. Over the next week the rearing temperature was increased to 10 °C and at 6 wph the fish were either challenged with *F. columnare* Fc7 (experimental group hereafter referred to as “Fc7”) or left unchallenged (“Ctrl"). Next, both challenged and unchallenged flasks were either treated with the antibiotic oxytetracycline (“AB”), the bacteriophage FCL-2 (“Phage”) or kept untreated (“None”), which resulted in the following six experimental groups: Ctrl_None, Ctrl_Phage, Ctrl_AB, Fc7_None, Fc7_Phage, Fc7_AB. Each condition was replicated in three replicate flasks yielding a total of 36 fish flask, containing 15 fish each (2 microbial states (germ-free/colonized) x 2 challenge states (Ctrl/Fc7) x 3 treatment (AB/Phage/None) x 3 biological replicates; Supp. Fig. 1). The mortality in each flask was assessed regularly until the experiment was terminated at 10 dpc. Additionally, samples were taken from the rearing water and from three fish per flask at 0 dpc (before challenge), 0.5, 2 and 10 dpc (see below) for microbiome analysis. Unfortunately, no sequencing data could be obtained for most of the fish samples, and samples from the fish microbiota are therefore not included in this analysis. Further, the bacterial density in the water was quantified by flow cytometry right before and after *F. columnare* was added to the samples (-0 and +0 dpc samples) and at 0.5, 1, 2, 4, 6, 8 and 10 dpc.

**Fig. 1:**
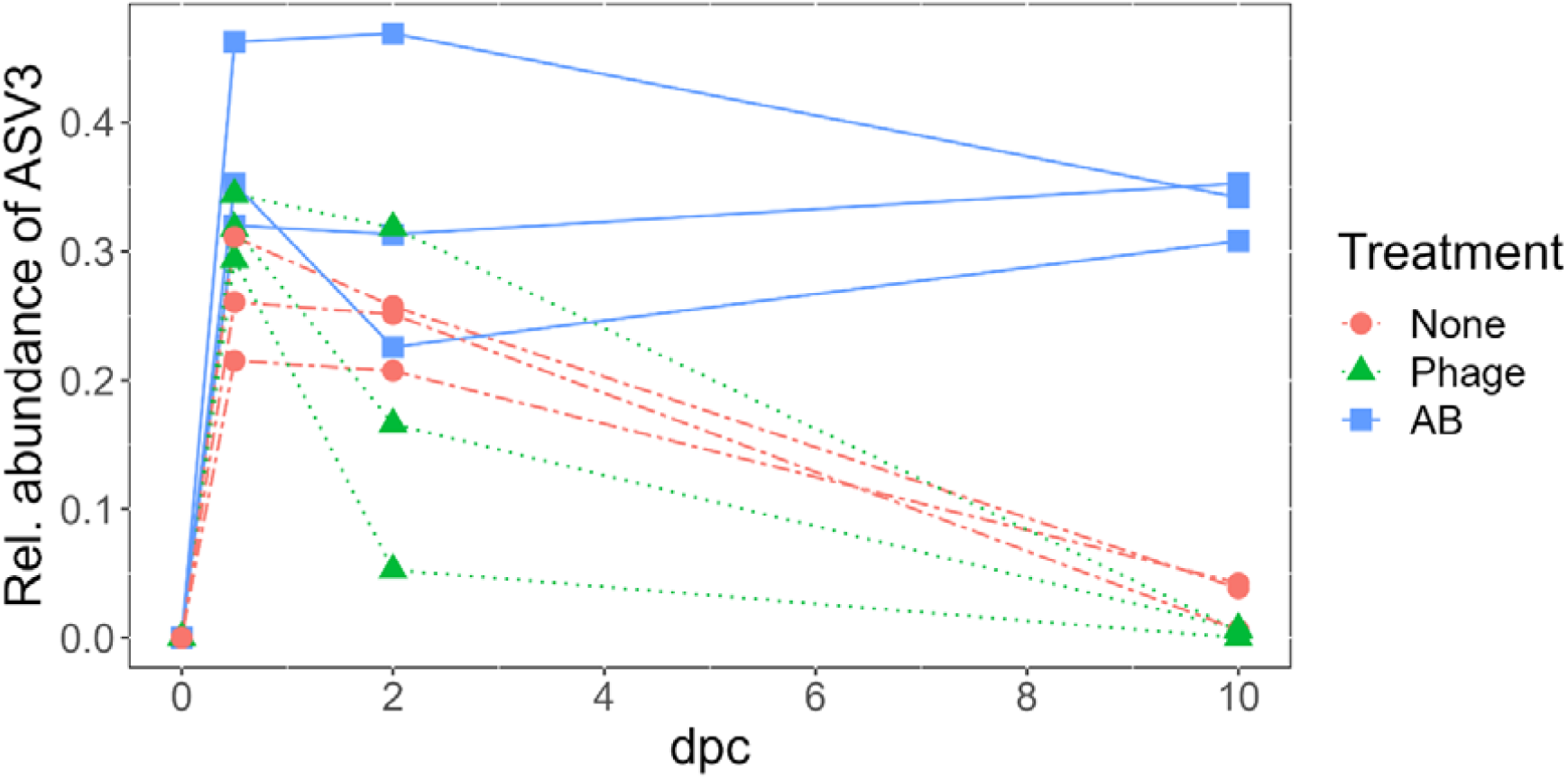
The relative abundance of ASV3 in the water microbiota of all Fc7-flasks of Exp.2. ASV3 was identified to represent *F. columnare* Fc7. Each line connects the observations within one rearing flask, and colours indicate the treatment application (None = control treatment, Phage = FCL-2 addition, AB = oxytetracycline addition).

Experiment 3 had the same setup as Exp.2, with the difference that the temperature was increased to 14 °C over the course of one week at 5 wph (Supp. Fig. 1). We increased the temperature in this experiment to 14 °C as we unexpectedly did not see mortality in the fish in Exp.2. Therefore, Exp.3 was conducted to investigate whether *F. columnare* Fc7 induced mortality at 14 °C, whether possessing a microbiota protects the fish against mortality and finally to evaluate whether phage-or antibiotic treatments impacts fish viability. No microbiota samples were taken from this experiment.

### Fish husbandry

We raised the Atlantic salmon as described by Gomez de la Torre Canny et al.^[41]^. In brief, Atlantic salmon eggs were received at ca. 78% development from AquaGen AS (Hemne, Norway) and immediately transferred to a dark room at 6 °C. Groups of 100 eggs were placed in a petri dish (13.5 cm Ø) and covered with Salmon Gnotobiotic Medium (SGM; 0.5 mM MgSO_4_, 0.054 mM KCl, 0.349 mM CaSO_4_ and 1.143 mM NaHCO_3_ dissolved in ultrapure water, autoclaved prior to use at 121 °C for 20 min). After one day, the eggs were sterilized (see below) and distributed into 250 ml cell culture flasks with a vented cap, containing 100 ml SGM and 17 fish eggs each. The eggs, and later the fish, were reared in these flasks until the end of the experiment. To maintain good water quality, 60% of the rearing water was exchanged 3 times a week. Dead fish were removed as soon as they were observed. For sampling and for terminating the experiment, fish were euthanised by a lethal dose of tricaine (20 mM, 0.2 µm sterile filtered). As the yolk sac fry that was used in this study is not considered as live animals under Norwegian legislation, the experiments conducted in this study had not to be approved by an animal welfare committee.

### Sterilization of fish eggs and reintroduction of bacteria (colonization)

The eggs were disinfected as described by Gomez de la Torre Canny and co-workers^[41]^. First, eggs were surface-sterilized 24 h after arrival at our laboratory by submerging them in an antibiotic cocktail (10 mgl^-1^ Rifampicin, 10 mgl^-1^ Erythromycin, 10 mgl^-1^ Kanamycin, 100 mgl^-1^ Ampicillin, 250 µgl^-1^ Amphotericin B, 150 mgl^-1^ Penicillin and 75 mgl^-1^ Oxolinic acid) for 24 h. Second, groups of 17 eggs were incubated in a Buffodine® solution (FishTech AS) containing 50 mgl^-1^ available iodine for 30 minutes. They were washed four times with 50 ml SGM and were then placed into a 250 ml cell culture flask containing 100 ml SGM.

To confirm axenity, sterility checks were performed on the hatching day, one week before challenge and at the end of the experiment. The sterility check was conducted for each germ-free flask by adding 100 µl rearing water to 3 ml of four different liquid media (Brain Heart Infusion, Glucose Yeast Broth, Sabourad-Dextrose Broth and Nutrient Broth) and a TSB agar plate. The four broths and the plates were incubated at room temperature (RT) for three weeks. If growth was observed in either medium, the flask was considered contaminated and removed from the experiment. In addition, water samples were taken and analysed for presence of bacteria using a flow cytometer (Attune NxT, ThermoFisher).

For generating microbially colonized flasks from germ-free flasks, bacteria were reintroduced at 1 wph by adding 60 ml of water from Lake Jonsvatnet (Trondheim, Norway) during the water change. The water was taken from a 50 m depth in February 2022 (Exp.2) and in June 2022 (Exp.3).

### *F. columnare* challenge and treatment of the fish with oxytetracycline or bacteriophage FCL-2

The number of fish was adjusted to 15 fish per flask at 5 wph and, depending on the experiment and experimental group, the temperature was steadily increased to either 9.8 ± 0.3 °C, 14.1 ± 0.3 °C or was kept at 5.6 ± 0.4 °C. For challenging the fish, *F. columnare* Fc7 (kindly provided by David Perez-Pascual and Jean-Marc Ghigo, Institute Pasteur, Paris;^[4]^) was grown in liquid TYES medium (0.5 gl^-1^ MgSO_4_ * 7 H_2_0, 0.2 gl^-1^ CaCl_2_ * 2 H_2_0, 0.4 gl^-1^ yeast extract, 4 gl^-1^ tryptone, 0.5 gl^-1^ D-glucose) at RT and 180 rpm overnight and harvested in late exponential phase at an OD_600_ of approximately 1. The bacterial culture was first spun down at 13.000xg for 10 min, then the pellet was washed with SGM once and finally resuspended in SGM resulting in a concentration of 1 x 10 ^9^ CFU m^-^l^1^. One ml was added per challenged fish flask, resulting in a theoretical final concentration of about 10 ^7^ *F. columnare* Fc7 CFUs ml^-1^ in the challenged fish flasks.

In Exp.2 and 3, treatment with either the phage FCL-2 against *F. columnare* Fc7 or the antibiotic oxytetracycline was applied immediately after challenge and daily the next ten days. For phage treatment we added 1 x 10^9^ PFUs per fish flask daily yielding an MOI (multiplicity of infection) of 1 at 0 dpc. For the antibiotic treatment, 4 mg of oxytetracycline (0.4g/l 0.2 µm sterile filtered stock) was added daily per flask, initially yielding 40 mgl ^-1^ per flask at 0 dpc. Nothing was added to the untreated control flasks.

### Preparation of phage stock

Phage strain FCL-2 ^[38]^ against *F. columnare* was kindly provided by Lotta-Riina Sundberg (University of Jyväskylä). Susceptibility of *F. columnare* Fc7 against FCL-2 was confirmed using the soft-agar overlay method. For that, *F. columnare* Fc7 was grown in liquid TYES medium at RT and 180 rpm under aerobic conditions until the exponential phase. Of this culture, 1 ml was added to 3 ml 50 °C warm soft TYES agar (TYES broth containing 7.5 gl^-1^ agar), vortexed and poured out on a TYES agar plate (containing 15 gl ^-1^ agar). The plate was incubated at RT for 1 h before 5 µl of the phage stock was added onto the plate. Formation of plaques indicated that *F. columnare* Fc7 was susceptible towards phage strain FCL-2. A phage stock was prepared by harvesting phages from soft TYES agar plates as propagation of the phage in liquid culture was not possible^[38]^. For that, 100 µl of phage solution (undefined concentration) were mixed with 3 ml soft TYES agar and 1 ml culture of strain Fc7 and was poured on a TYES agar plate, which was incubated at RT overnight. The soft top agar containing the phages was scraped off and suspended in sterile SM buffer (5.8 gl^-1^ NaCl, 50 mll^-1^ Tris buffer (1 M, pH 7.5), 2 gl ^-1^ MgSO_4_ * 7 H_2_0) at a volume ratio of 1:1 soft agar to SM buffer. The mixture was vortexed, spun down at 5,000xg for 10 min and filtered through a 0.2 µm filter. The phage stock was concentrated by first centrifuging at 22,000xg for at least 8 h, before the supernatant was removed and the phage pellet resuspended in a small volume of SM buffer. The titer of the phage stock was determined by spotting out serial dilutions of the stock on soft-agar-overlaid plates and counting plaques.

### Flow cytometry analysis

We quantified the bacterial density in the rearing water in Exp.2 using flow cytometry at nine different timepoints: right before and after *F. columnare* Fc7 was added to the samples (-0 and +0 dpc samples) and at 0.5, 1, 2, 4, 6, 8 and 10 dpc. We sampled 1 ml water per flask and sampling time. The water samples were fixated in 0.1% glutaraldehyde for 15 min before they were snap-frozen in liquid nitrogen and stored at -80 °C until data acquisition. Samples were diluted in 0.2 µm-filtered phosphate-buffered saline to obtain stable sample aquation and dilute background noise present in the sample. The water samples were stained with the DNA-binding fluorescent dye SYBR green I (Invitrogen, final concentration 2x), vortexed and incubated for 15 minutes in the dark at 37° C. Stained samples were vortexed and 160 µl was sampled at a 100 µlmin ^-1^ flowrate. A zip clean of the instrument was performed between approximately every 6^th^ sample. Data were collected using the blue laser (488 nm) with detection in BL1 (530/30 nm) and BL3 (695/40 nm) using a BL1 threshold of 1500-3000 (depending on sample). Instrument voltages were as follows; FSC 320V, SSC 260V, BL1 320 V and BL3 350V. Filtered PBS and 0.2 µm-filtered fish rearing water were used as negative controls and a pure culture of strain Fc7 as positive control. Identification of bacterial populations was achieved by comparing 0.2 µm-filtered fish rearing water with unfiltered rearing water. All samples were gated in the same way with some minor modifications to each sample gate to ensure high quality data (figshare.com/articles/dataset/fsc_files_Fc7_Salmon/21518922/1).

### Sampling for characterization of the water microbiota in Exp.2

Water samples for microbiome analysis were taken only for Exp.2. Samples were taken at four timepoints and only from colonized flasks: Right before challenge with *F. columnare* Fc7 (0 dpc) and at 0.5, 2 and 10 dpc. For each sampling time we collected the rearing water bacterial community by filtering 10 ml rearing water through a 0.2 µm polycarbonate filter (Osmonics, 25 mm diameter). The filter was cut in pieces and transferred to a BeadBashing tube of the ZymoBIOMICS^TM^ 96 MagBead DNA kit (Zymo), snap-frozen in liquid nitrogen and stored at -80°C. In total 72 water samples were taken.

### DNA extraction

DNA was extracted from all water samples using the ZymoBIOMICS ^TM^ 96 MagBead DNA kit (Zymo) and a KingFisher Flex instrument. The filters representing the water samples were cut in smaller pieces using a sterile scalpel prior to extraction of DNA. The filters were homogenized by adding 750 µl (450 µl for 0 dpc samples) lysis buffer from the DNA extraction kit and run two cycles à 30 seconds at 5500 rpm in a Precellys 24 (Bertin Technologies). The homogenized samples were spun down at 13,000xg for 10 min. For samples from 0 dpc, we used 450 µl of the homogenate supernatant, whereas 200 µl were used from the 0.5, 2 and 10 dpc samples. DNA was extracted following the manufacturer’s protocol except for final elution in 100 µl DNAse-free water (50 µl for 0 dpc samples). The extracted DNA was stored at -20 °C until library preparation. Additionally, two negative controls using DNA-free water as input for DNA extraction were prepared.

### 16S rRNA amplicon sequencing

The v3+v4 region of the 16S rRNA gene was amplified using the primers Ill-341F (5’- TCGTCGGCAGCGTCAGATGTGTATAAGAGACAGNNNN**<u>CCTACGGGNGGCWGCAG</u>**-3’) and Ill-805R (5’-GTCTCGTGGGCTCGGAGATGTGTATAAGAGACAGNNNN**<u>GACTACNVGGGTATCTAAKCC</u>**-3’), with the target sequences shown in bold^[42,43]^. PCR was conducted in 25 µl reaction volumes, where each reaction contained 0.15 µM of each primer, 0.25 µM of each dNTP as well as 0.4 U Phusion hot start polymerase and the respective buffer from Thermo Scientific. For samples from 0 dpc, 1 µl of a 1:100 dilution of the DNA extract was used as template, whereas 1 µl undiluted DNA extract was used as template for the other samples. The cycling conditions were as follows: an initial denaturation step at 98 °C for 60 seconds followed by 33 cycles of 98 °C for 15 sec, 55 °C for 20 sec and 72 °C for 20 sec. The final elongation step was 72 °C for 5 minutes before the samples were cooled to 10 °C. PCR products were evaluated by electrophoresis on a 1% agarose gels containing 50 µM GelRed (Biotium) for 1 h at 110 V. The samples were normalized using Sequal Prep^TM^ Normalization plates (96 wells, Invitrogen) and indexed using the Nextera® XT Index Kit v2 Sets A and D. For indexing, 2.5 µl normalized PCR product was used as template with 2.5 µl of each indexing primer, 0.2 µM of each dNTP, 0.4 U Phusion hot start polymerase and its buffer from Thermo Scientific (total volume 25 µl). The same cycling program as above was run with 12 cycles. The indexed PCR products were again normalized by loading 15 µl indexed PCR products onto Sequal Prep ^TM^ Normalization plates and were pooled and concentrated using a Amicon® Ultra 0.5 ml centrifugal filter (30K membrane, Merck Millipore). A NanoDrop™ One Microvolume Spectrophotometer (Thermo Scientific™) was used to evaluate the quality and quantity of the DNA. The samples were sent to the Norwegian Sequencing Center using one run on a MiSeq v3 instrument with 300 paired ends. The sequencing data was deposited at the European Nucleotide Archive (ERS14896569-ERS14896640)

### Analysis of the Illumina sequencing data

The USEARCH pipeline (v.11) ^[44]^ was used to process the data obtained from Illumina sequencing. The sequencing pairs were merged, and primer sequences trimmed off using the Fastq_mergepairs command with a minimal length of 390 bp. The merged sequences were quality-filtered using the Fastq-filter function with the default error threshold value of 1. The reads were pooled, dereplicated and singleton reads removed. Amplicon sequence variants (ASVs) were generated using the Unoise2 command ^[45]^ with the default minimum abundance threshold of 8 reads in the total dataset. Taxonomical assignment of the ASVs was achieved using the Sintax command ^[46]^ with a confidence threshold of 0.8 and the RDP reference dataset v. 18 ^[47]^. Reads classified as eukaryotes or chloroplasts were removed from the data set. A few ASVs that were highly abundant in negative controls for the DNA extraction or the phage stock, but less abundant in the samples were considered to represent contaminating DNA and were removed from the data set. One sample (replicate flask 2, 0.5 dpc, Ctrl-Phage group) was removed from the dataset as it had extremely poor sequencing efficiency compared to all other samples. In the final ASV table, samples had 108,719 ± 20,015 reads on average and were normalized by scaling to 74,545 reads per sample. All analyses were performed using the normalized ASV table. By using the BLAST algorithm^[48]^ the 16S rDNA sequence of *F. columnare* Fc7 (Supp. Fig. 2) was compared to all ASVs of the dataset and we identified ASV3 to correspond to *F. columnare* Fc7.

**Fig. 2:**
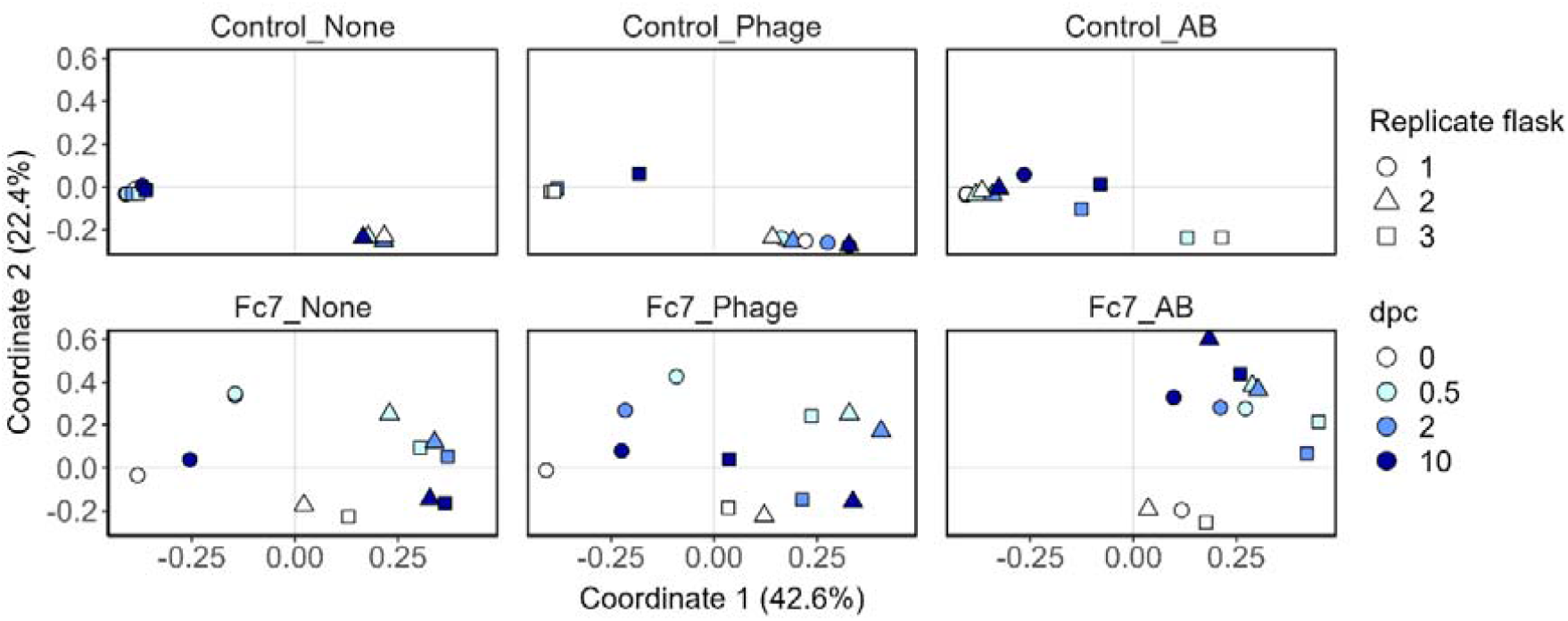
PCoA of the Bray-Curtis dissimilarities of all samples from Exp.2. Sampling timepoints are represented by different colours, whereas biological replicate flasks are indicated by different shapes. All panels are from the same PCoA but were facetted into the different treatment groups.

### Statistical analysis

All statistical analyses were performed in R (v. 4.1.3 ^[49]^) using RStudio (2022.07.1) and the packages Phyloseq (v. 1.38.0 ^[50]^), Vegan (v. 2.6.2 ^[51]^), ggplot2 (v. 3.3.6), dplyr (v. 1.0.9), reshape2 (v. 1.4.4), genefilter (1.76.0), DECIPHER (v. 2.22.0) and ggh4x (v. 0.2.2.9000). The *renyi* function of Vegan was used to calculate the α-diversities of samples as Hill’s diversity numbers^[52,53]^. Ordination by principal coordinate analysis (PcoA) was performed using the *ordinate* function from phyloseq for Bray-Curtis dissimilarities, if not stated otherwise. Phylogenetic trees were generated using the phangorn package in R (v. 2.9.0^[54]^) by first constructing a neighbour-joining tree and then fitting a GTR+G+I model to it. PERMANOVA analyses^[55]^ based on Bray-Curtis dissimilarities (if not stated otherwise) were performed using the *adonis*2 function from vegan by running it in 100 iterations with 999 permutations each and the mean p-value of the 100 iterations was reported (mathematically lowest possible p-value = 0.001). For statistical univariate data (e.g. α-diversity indices or abundance of certain ASVs), the data was checked for normality using the Shapiro-Wilk test (*11hapiro.test* function). When the data were found to be normally distributed, a Welch’s t-test was used for data with two groups and ANOVA for data with three groups. A Mann-Whitney U test or Kruskal-Wallis test was performed for these purposes on non-normally distributed data. Significant ANOVA or Kruskal-Wallis tests were followed by a Bonferroni-corrected Dunn’s test. A significance level of α < 0.05 was used for all analyses. All box plots are presented as median and upper and lower quartile as box, whiskers include all samples except for outliers. Kaplan-Meier survival analysis was performed using the survival (v.3.3.1) and survminer (v. 0.4.9) packages in R.

## Results

### Challenging Atlantic salmon yolk sac fry with *Flavobacterium columnare* Fc7 at different temperatures

In the first experiment (Exp.1), we examined whether the bacterial pathogen *Flavobacterium columnare* Fc7 induced a lethal infection in germ-free Atlantic salmon yolk sac fry at 6, 10 or 14 °C. Both at 10 and 14 °C, all fish died within 60 h and 48 h, respectively, in all three replicate flasks that had been challenged with *F. columnare Fc7* (data not shown). No mortality was observed in flasks challenged at 6 °C or in any of the control flasks that had not been added *F. columnare* Fc7. The exception was one dead fish in a flask that was exposed to the bacterial commensal *Janthinobacterium* sp. 3.108 at 14 °C. We therefore concluded that *F. columnare* Fc7 induced mortality in germ-free Atlantic salmon yolk sac fry at both 10 and 14 °C.

### *F. columnare* Fc7 challenge and phage treatment at 10°C in Exp.2

As yolk sac fry thrive best at lower temperatures, we performed the second challenge experiment (Exp.2) at 10 °C. We challenged both germ-free and microbially colonized fish with *F. columnare* Fc7 to assess whether the presence of a microbiota protects the fish against lethal infections. By consecutively treating them with either the bacteriophage FCL-2 or the antibiotic oxytetracycline (OTC) we wanted to examine whether bacteriophage treatment can be used to protect the fish against infection and further assessed the effect of phage-and antibiotic treatment on the water and fish microbiota by 16S rDNA sequencing (see Supp. Fig. 1 and Material and Methods).

#### Effect of phage therapy on fish survival at 10 °C

Unexpectedly, *F. columnare* Fc7 did not induce mortality in neither the germ-free nor the colonized fish in Experiment 2. Thus, we could not reproduce the observed mortality of Exp.1. Therefore, we could not draw a conclusion whether phage treatment or presence of a microbiota protected the fish against lethal infections in Exp.2.

#### Effect of phage FCL-2 on the relative amounts of F. columnare Fc7 in the water microbiota

We sampled rearing water and fish from Exp.2 at 0, 0.5, 2 and 10 days post challenge (dpc) in order to investigate the impact of phage-and antibiotic treatment on the water and fish microbiota by 16S rRNA gene sequencing. Unfortunately, the amplification of bacterial 16S rDNA from most fish samples was unsuccessful, and therefore, only the bacterial community of the water samples is presented.

We first investigated whether phage therapy reduced the relative abundance of the pathogen in the rearing water. We did not observe the ASV representing *F. columnare* Fc7 (ASV3) in the water prior to the *F. columnare* Fc7 challenge (0 dpc samples; Fig. 1). In samples collected at 0.5 dpc, the relative average abundance of ASV3 in the bacterial communities of the water was 31 ± 7 % in the flasks to which strain Fc7 was added (Fc7-flasks; Fig. 1). There was no difference in the relative abundance of ASV3 at 0.5 dpc among the Fc7-flasks (Kruskal-Wallis test, p = 0.433), indicating that similar amounts of *F. columnare* Fc7 were added to each flask. The relative abundance of ASV3 decreased in both the Fc7_None and the Fc7_Phage flasks. This decrease was on average 76-fold from 0.5 to 10 dpc in Fc7_Phage flasks but only 9-fold in Fc7_None flasks (Fig. 1). Due to the limited numbers of replicates, no statistical test could be conducted to confirm whether the difference between treatments were significant. Further, the relative abundance of *F. columnare* Fc7 decreased faster in most Fc7_Phage flasks than in the Fc7_None flasks (Fig. 1), indicating that phage therapy successfully reduced the populations of *F. columnare* Fc7 in the flasks.

Unexpectedly, the antibiotic treatment did not affect the relative abundance of ASV3 in the water community profiles (Fig. 1). Moreover, the relative abundance of ASV3 in Fc7_AB flasks was significantly higher than in Fc7_None and Fc7_Phage flasks at 10 dpc (Kruskal-Wallis test, p = 0.049). This was surprising, as preliminary tests confirmed susceptibility of *F. columnare* Fc7 to OTC on TYES agar plates (data not shown).

In conclusion, the relative abundance of *F. columnare* Fc7 decreased in both Fc7-None and Fc7-Phage flasks, with a slightly stronger decrease in Fc7-Phage flasks, while no decrease was observed in Fc7-AB flasks.

#### Impact of F. columnare Fc7 challenge on the rearing water microbiota

The water microbiota differed significantly between all flasks already at 0 dpc, prior to challenge with *F. columnare* Fc7 even though all flasks were treated identically until the challenge with *F. columnare* Fc7 (Fig. 2 and Fig. 3; PERMANOVA: p-value = 0.001). These differences were therefore most likely a result of individual bacterial community succession in each flask during the six weeks before the challenge and not of the experimental groups.

**Fig. 3:**
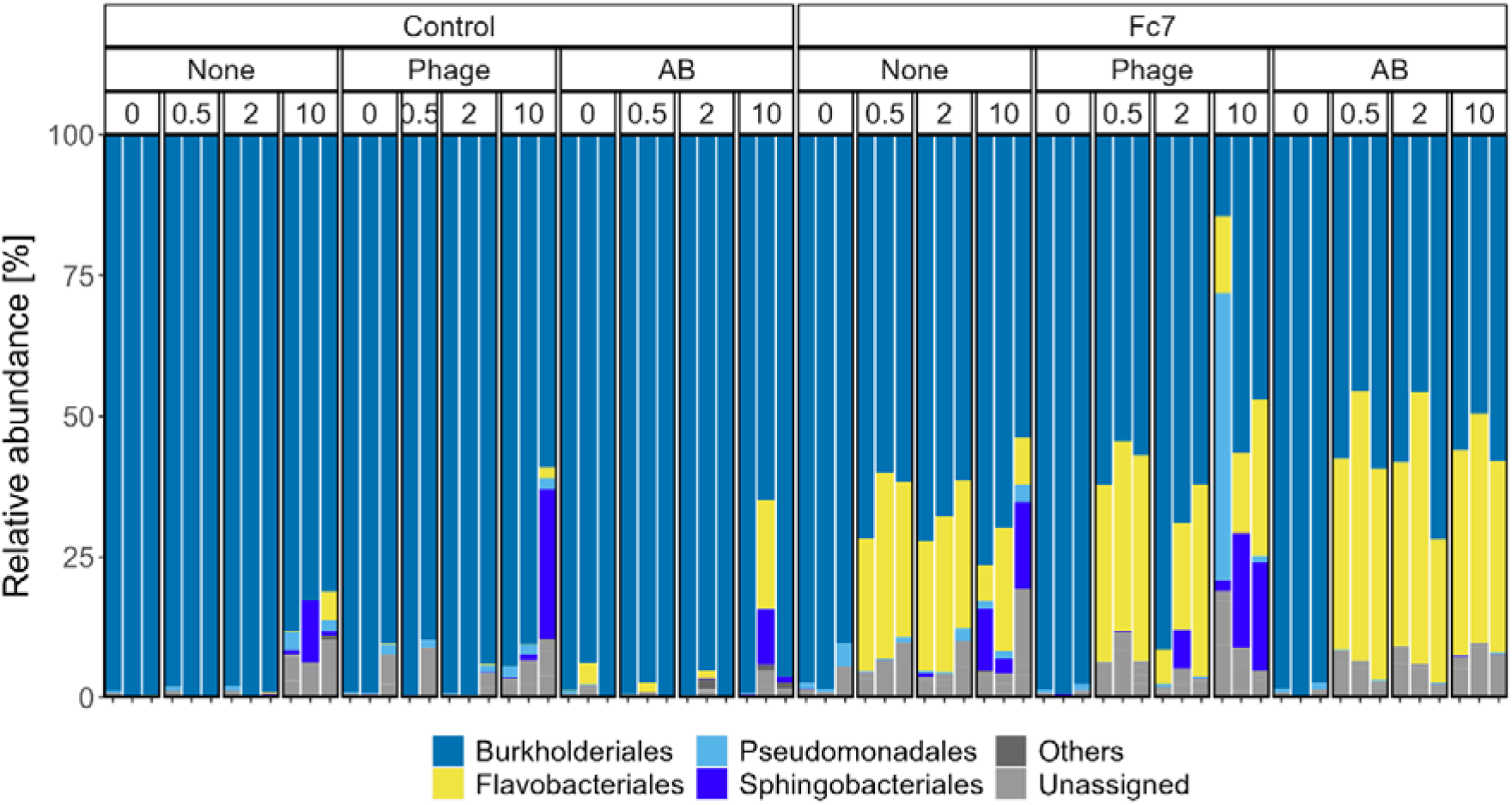
Bacterial community composition of each water sample from Exp.2, shown at the order level. Orders that are not appearing with at least 5% in at least one sample are summarized as “others”.

The microbiota in the Ctrl-flasks (no *F. columnare* Fc7 added) did not change significantly during the experiment (PERMANOVA, p > 0.580 for all three comparisons; Fig. 2). However, in the Fc7-flasks, the water microbiota had changed 12 hours after addition of strain Fc7, mainly due to an increase in the relative abundance of ASV3 (Fig. 2, 3 and Supp. Fig. 3). Interestingly, PCoA indicated that 10 days after addition of *F. columnare* Fc7, the water microbiota in Fc7-Phage and Fc7-None flasks returned to their composition prior to the addition of *F. columnare* Fc7 at 0 dpc (Fig. 2). This recovery was, however, not observed in Fc7_AB flasks (Fig. 2), as the relative abundance of ASV3 remained high (Fig.1 and Supp. Fig. 3).

These temporal effects in the Fc7-flasks were also observed when ASV3 was removed from the dataset prior to PCoA ordination (Supp. Fig. 4) and when ordinations were based on presence-absence data (Sørensen-Dice dissimilarity, Supp. Fig. 5). These findings show that addition of *F. columnare* Fc7 affected the water microbiota in the challenged flasks.

**Fig. 4:**
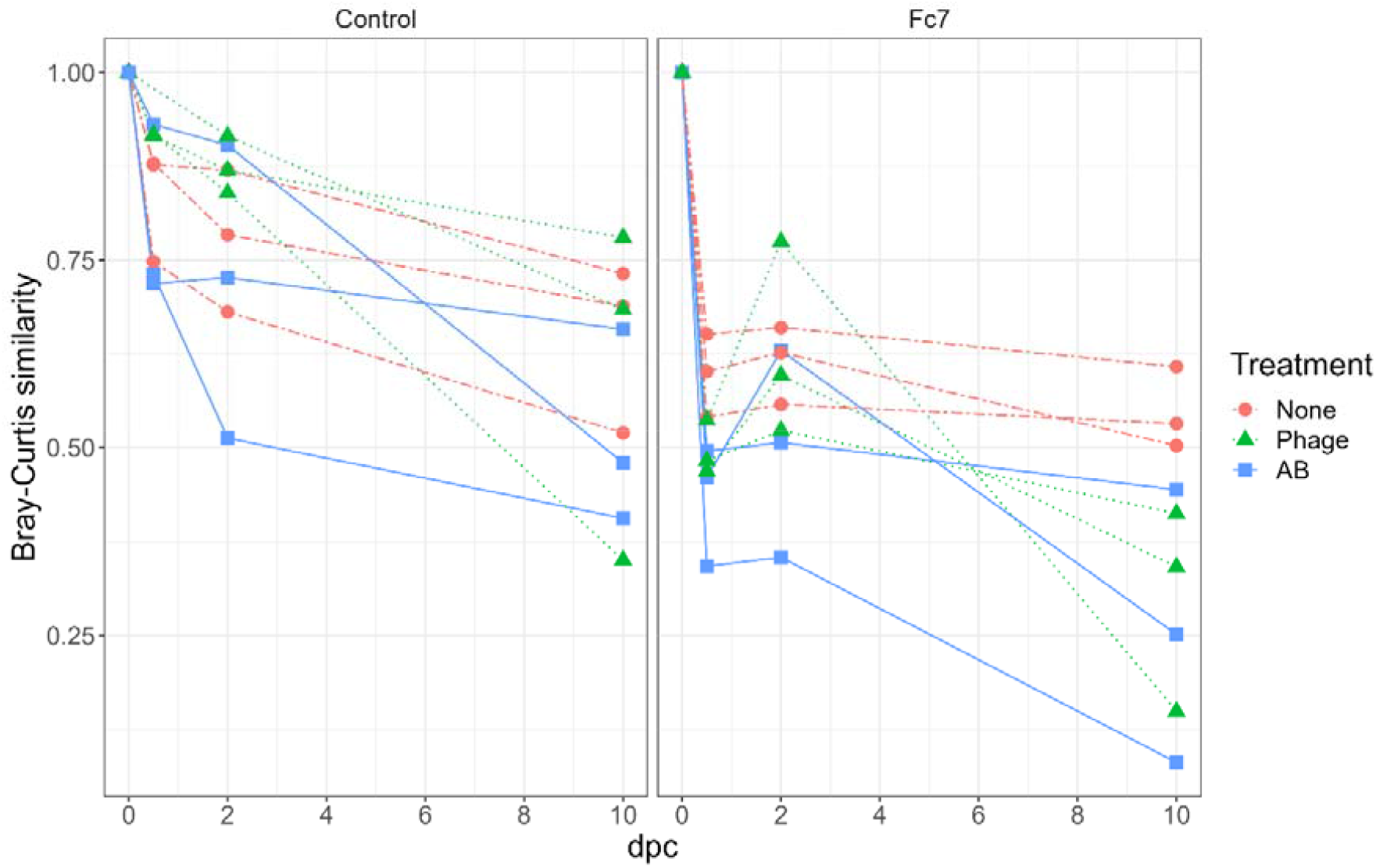
Bray-Curtis similarities within each replicate flask comparing the water bacterial community from each timepoint to the community at 0 dpc. Each line represents one rearing flask, and colours indicate the treatment application (None = control treatment, Phage = FCL-2 addition, AB = oxytetracycline addition).

#### Effect of phage therapy and antibiotic treatment on the water microbiota

To evaluate the effect of the treatments on the bacterial communities we evaluated the community composition change over time, the α-diversity and the bacterial density.

We therefore compared the bacterial community in the None-flasks with Phage-or AB-flasks over the course of the experiment to assess the effect of phage-and antibiotic treatment both in the presence and absence of *F. columnare* Fc7. Because the bacterial communities differed between replicate flasks from the same experimental group (Supp. Fig. 6) we investigated the temporal change within each individual flask. For each flask, we calculated the Bray-Curtis similarity between each sampling timepoint and the start of the experiment (0 dpc). For Ctrl-flasks, the change in B ray-Curtis similarity over time was comparable between treatments (Fig. 4). This indicated that in the absence of *F. columnare* Fc7, neither the antibiotic-nor the page treatment influenced the water bacterial community. For the Fc7-flasks, the communities in Fc7_Phage and Fc7_AB flasks changed more over time than Fc7_None flasks (Fig. 4). Therefore, in the presence of the pathogen, both the antibiotic and the phage treatment appeared to influence the bacterial water communities compared to Fc7_None flasks.

Hill’s diversity of order 1 (^1^D; exponential Shannon index) was not significantly affected by the phage treatment (Supp. Fig. 7), as no significant difference was observed when comparing the change in^1^D from 0 to 10 dpc for each flask between the different treatments (ANOVA, p = 0.532 and 0.592 for Ctrl-and Fc7-flasks, respectively). Highly unexpectedly, AB treatment increased the α-diversity in the Ctrl-flasks by a factor of 2 from 0 to 10 dpc, whereas this was not observed in Ctrl-None flasks (Supp. Fig. 7). Lastly, phage treatment did not decrease the bacterial density when Fc7 was absent (Supp. Fig. 8).

These findings indicate that phage therapy did not influence the bacterial communities in the absence of the phage’s host, *F. columnare* Fc7, whereas changes in the microbiota were observed in the presence of the phage’s host. Phage therapy did further not significantly affect the α-diversity. Unexpectedly, also the antibiotic oxytetracycline did not disturb the microbiota in the water in the absence of *F. columnare* Fc7.

### *F. columnare* Fc7 challenge and phage treatment at 14 °C in Exp.3

As no mortality was induced by *F. columnare* Fc7 at 10 °C in Exp.2, we conducted a third experiment to investigate the protective effect of possessing a microbiota and to determine whether phage and antibiotic treatment reduced mortality in infected fish. The experimental setup was similar between Exp.2 and 3, with the exception that the temperature was increased to 14 °C to increase infectivity *of F. columnare* Fc7^[35]^.

All germ-free fish in Fc7_None flasks died after challenge with *F. columnare* Fc7. In contrast, mortality was only observed in one flask in colonized Fc7_None flasks (Fig. 5). Thus, the survival was significantly higher when fish possessed a microbiota (Kaplan-Meier survival analysis, p<0.0001). This suggests that the fish microbiota protects the fish against lethal bacterial infection.

**Fig. 5:**
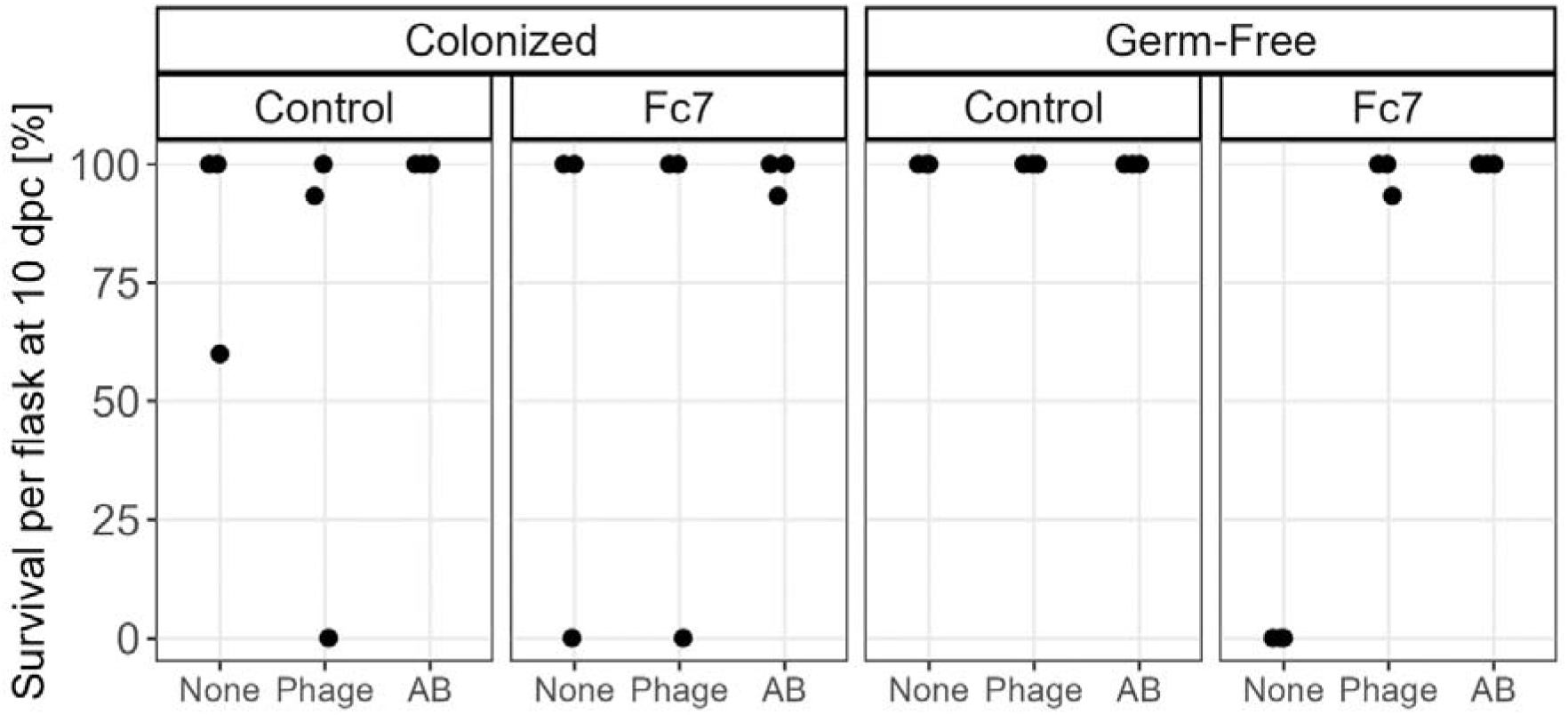
Fish survival at 10 dpc in Exp.3. Each dot represents survival in one replicate flask (three replicate flasks per group).

As there were no mortality in neither phage-nor the antibiotic-treated germ-free flasks, we concluded that both phage therapy and antibiotic treatment protected the fish against lethal infections (Fig. 5). However, due to high survival in colonized fish we could not conclude on the effect of phage therapy in colonized fish. We further observed unexpected mortality in colonized Ctrl-None and Ctrl-Phage flasks to which no pathogen was added. These observations are addressed in the Discussion part.

We therefore concluded that the presence of a microbiota protected the yolk sac fry against lethal infection with *F. columnare* Fc7, and also that both the phage-and the antibiotic treatment protected the fish.

## Discussion

We urgently need alternatives to antibiotics as antibiotic resistance genes spread, the development of novel antibiotics is slow and the ones we use have harmful side effects on our microbiome^[10,56]^. One alternative is phage therapy, where a virulent bacteriophage is used to lyse, and thus kill, a specific pathogenic bacterium ^[26]^. Clinical trials on humans with phage therapy were shown to be challenging due to administrational hurdles and due to a lack of information on how phages affect the human body and its microbiome^[57,58]^.

In aquaculture, several commercial phage-products are already available^[27]^. The application of phage therapy often increases survival in infection experiments (e.g.^[59–61]^), although sometimes studies find no benefit from the therapy ^[62]^. While it is generally accepted that phage therapy may alleviate aquatic disease outbreaks, the effect that phage therapy has on the microbiota is, however, poorly documented. In this study, we therefore aimed to investigate the impact of phage therapy on the rearing water of the commercially important species Atlantic salmon.

In order to test phage therapy, we first had to establish a challenge protocol to infect the fish with a bacterial pathogen, and we chose to use *F. columnare* for this purpose. In Exp.1, we found that *F. columnare* rapidly induced mortality in Atlantic salmon yolk sac fry at both 10 and 14 °C. This observation could be replicated at 14 °C in Exp.3 but no at 10 °C in Exp.2. It was puzzling that we observed mortality in germ-free fish at 10 °C in Exp.1 but not in Exp.2 as the experimental conditions were identical. Differences between egg batches might have contributed, although it is impossible to investigate this in retrospect. Because *F. columnare* primarily infects fish at warmer temperature, we expected less infection success at lower temperature^[35,36]^. Still, it was interesting that infection induced mortality abruptly, with only a 4 °C difference between the experiments.

After establishing a challenge protocol, we first investigated whether the microbiota in the fish has a protective effect during the bacterial challenge and therefore compared fish survival in germ-free and colonized fish. All germ-free fish infected with *F. columnare* at 14 °C died within three days. In contrast, for the colonized fish, mortality was only observed in one of the three replicate flasks. Thus, the microbiota of the salmon protects the fish. Such a protective effect of the microbiota is generally assumed to be present in animals, however, studies investigating it in fish are scarce so far^[4,5]^. It was therefore important to confirm this protective effect in fish, after it had been shown previously in zebrafish^[5]^ and rainbow trout^[4]^.

The high mortality in untreated germ-free fish, and the fact that we saw no mortality in phage-treated germ-free fish, showed that bacteriophage therapy was successful in increasing survival during infection with *F. columnare*. The same phage that we used (FCL-2) has also been successfully used earlier in rainbow trout and zebrafish, where elevated survival also was observed ^[38,40]^. Our findings therefore confirm the potential of using this phage in aquaculture against *F. columnare* infections.

Unfortunately, we were not able to draw a definite conclusion about the success of phage therapy in the colonized fish, as survival in the colonized Fc7-flasks was generally very high. Unexpectedly, we occasionally observed that fish suddenly died, even in flasks that were unchallenged with *F. columnare* Fc7. This was likely caused by instabilities in our experimental system at 14 °C: At higher temperature, increased bacterial growth and metabolic activity may deplete oxygen, causing subsequent fish death. This temperature-dependent mortality has been observed previously in similar experimental setups in our research group (data not shown). This explanation seems plausible, as no mortality was observed when OTC was added to the water. OTC decreases metabolic activity by inhibiting protein synthesis. Thus, the bacterial oxygen consumption could not increase when OTC was present in the water, and the fish did not die.

In order to assess the effect of phage therapy on the water bacterial community, we sampled the water microbiota in Exp.2. We originally aimed to investigate both the fish and the water microbiota. However, for over 70% of the fish samples, we could not generate 16S rRNA gene amplicons and, consequently, all fish samples were discarded from further analysis. Nevertheless, it is relevant to investigate the effect of phage treatment on the water bacterial community, as the water influences the fish microbiota^[63–66]^. First, we inspected whether phage treatment reduced the relative abundance of *F. columnare* in the water. Here, we expected that the phage strongly reduced the relative abundance of the pathogen due to lysis of the cells. However, the relative abundance of *F. columnare* Fc7 in the water microbiota was quickly reduced, also in flasks that were not treated after the challenge. This shows that *F. columnare* were not able to persist in the rearing flasks under the experimental conditions used in this study. *F. columnare* Fc7 might not be metabolically active at 10 °C ^[35]^, and this might decrease the effect of the phage treatment ^[67,68]^. Still, we observed a faster and more pronounced decrease of the relative *F. columnare* F c 7 abundance due to addition of phage FCL-2, which indicates that the phages were lysing the bacterial cells.

Phage therapy has been proposed to exert minor effects on the microbiota because the treatment only targets one specific population ^[69]^. When the phage’s target bacterium is absent, no strong effects on the microbiota are expected ^[70–73]^. Concordantly, we observed no significant changes in the community composition, α-diversity or bacterial density due to the FCL-2 treatment in the absence of *F. columnare*. When the phage’s target is present, however, the removal of the target bacterial population can affect the community properties^[31,33]^. These effects depend on the interaction network of the target population within the community and its abun d *F*a.n*c*c*o*e*lu*.*mnare* Fc7 was not present in the water communities prior to the Fc7 challenge and is therefore most likely not part of the interaction network in the microbial communities. However, we added *F. columnare* Fc7 in high abundances to the Fc7_Phage flasks. It was therefore not surprising to observe changes in the bacterial compositions in these flasks. These small changes might be due to lysis of Fc7-cells which liberates nutrients to the other community members ^[31]^. It would have been interesting to investigate the effect of phage therapy on the microbiota during a natural outbreak of columnaris disease. In such a scenario the removal of the pathogenic population might result in stronger downstream effects due to disruption of the bacterial interaction-network^[31]^. Even though slight changes in the bacterial community occurred, no effect on the α-diversity or absolute abundance of the cells was observed. With this, our study adds to the growing body of evidence, that phage therapy does not cause negative side-effects on the bacterial community^[30,59,74–76]^.

Surprisingly, antibiotic treatment with OTC did not affect the microbiota as we would have expected it. The antibiotic treatment did not reduce the relative abundance of *F. columnare* Fc7 and did further not distort the microbiota in the flasks or decrease α-diversity. This was unexpected, as OTC is a widely-used broad-spectrum antibiotic that has been shown to elicit strong disturbances onto the microbiota and to decrease the bacterial richness^[19,20,77–79]^. It is also generally accepted that antibiotic treatment in general has a disruptive effect on the microbiota^[80–82]^. Nevertheless, some studies find that OTC does not always disturb the microbiota or decrease the richness ^[83,84]^. A reason for why we did not see stronger effects on the bacterial community and the abundance of *F. columnare* Fc7 could be that OTC’s function was somehow impaired. However, its efficacy against *F. columnare* Fc7 was tested before, during and after the experiment and a fresh AB stock was prepared at 5 dpc in order to avoid degradation of OTC over time. Furthermore, the higher survival of the antibiotic-treated fish in Exp.3 indicated that OTC was active and functioning. Another explanation could be that the bacteria in the water are slow growing at 10 °C, which could delay observable effects by the bacteriostatic OTC. Since we unexpectedly did not observe disturbances induced by OTC, it would had been beneficial to use a bactericidal antibiotic for disturbing the bacterial communities, which would have allowed better comparison of the effects of antibiotics and phage therapy on the fish microbiota.

In conclusion we showed that phage therapy protected Atlantic salmon yolk sac fry against infection with *F. columnare*, without disturbing the microbiota of the water. While our work was conducted in Atlantic salmon, we think that our findings can also be useful for discussing usage of phage therapy in a context outside of aquaculture.

## Funding

MSG and AWF were supported by a PhD scholarship from NTNU, Faculty of Natural Science.

## Acknowledgements

We would like to thank Amalie Mathisen for valuable help in the lab. Further, we thank Rolf Myklebust from Aquagen AS for providing us with salmon eggs. We also thank Lotta-Riina Sundberg for providing us with phage FCL-2 and helpful tips on phage cultivation, as well as Jean-Marc Ghigo and David Perez-Pascual for providing us with *F. columnare* strain Fc7.

## Author contribution

AF, MG, EA, OV and IB designed and conceived the experiment. AF, MG and TV conducted the experiments and AF and MG analysed the data. AF wrote a first draft of the manuscript, and all authors improved and approved it.

## Conflicts of interests

The authors declare no competing interests.

## Data availability

All flow cytometry files used for bacterial density quantification are available through https://figshare.com/projects/fsc_files/152463. The Illumina sequencing reads are available through the European Nucleotide Archive (ERS14896569-ERS14896640).

## Notes

### Competing Interest Statement

The authors have declared no competing interest.

