## Supplementary Files for "Phage therapy minimally affects the water microbiota in an Atlantic salmon (*Salmo salar*) rearing system while still preventing infection"

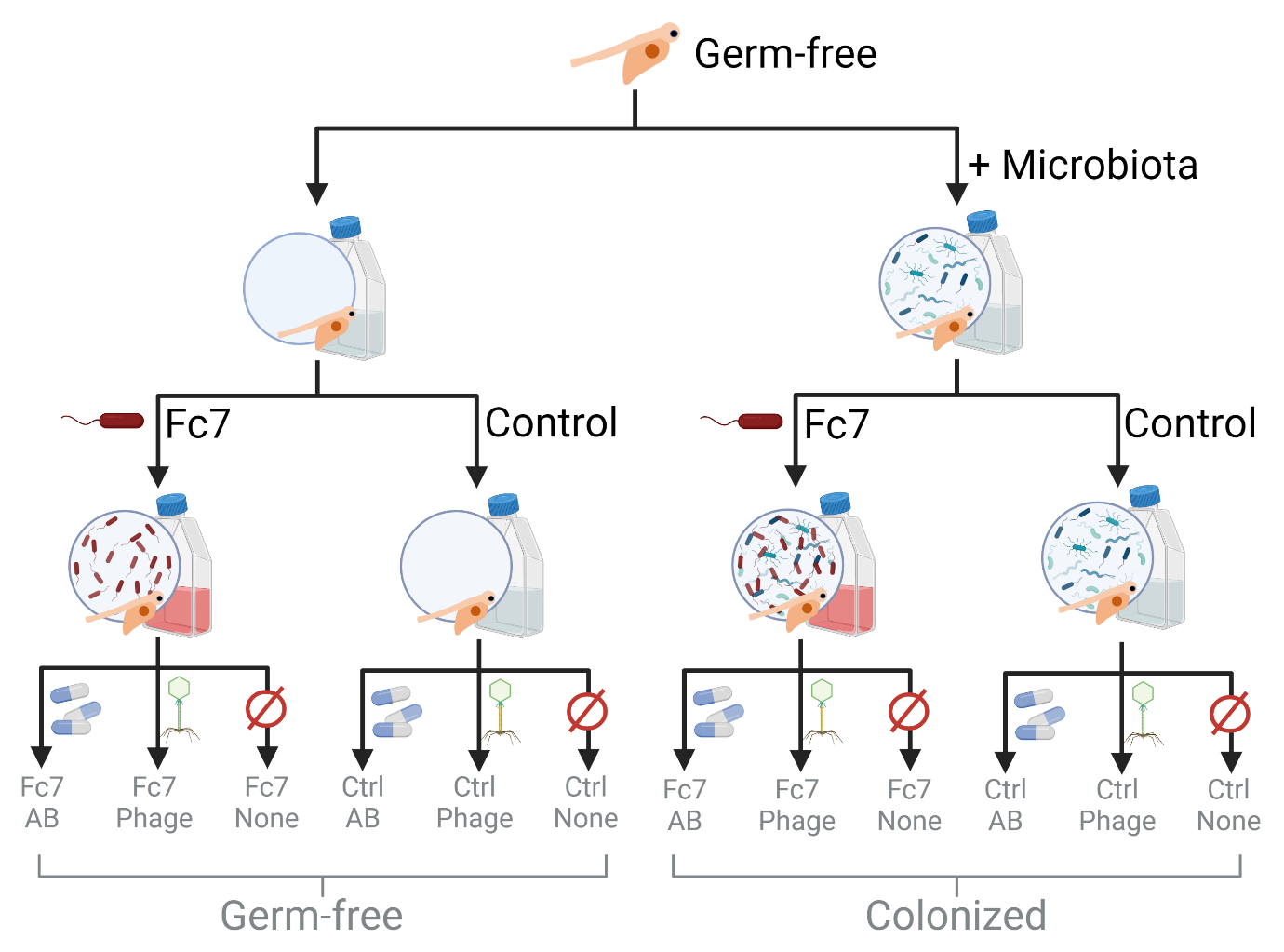

Supp. Fig. 1: Experimental design of Exp.2 and Exp.3, resulting in twelve experimental groups each. Each group consisted of three replicate flasks (36 in total), which each contained 15 fish. Antibiotics and phages were added daily throughout the 10 days following infection at 6 weeks post hatching (wph). Figure created with BioRender.com.

5’-TTAACACATGCAAGTCGAGGGGTAGAAGGAGCTTGCTCCTTTGAGACCGGCGCACGGGTGCGTAACGCGTATGCAATCTACCTTGTACAGGGGGATAGCCCAGAGAAATTTGGATTAATACCCCATAGTATTTTCAGATGGCCTCATTTGATTATTAAAGTTCCAACGGTACAAGATGAGCATGCGTCCCATTAGCTAGTTGGTGTGGTAACGGCATACCAAGGCAACGATGGGTAGGGGTCCTGAGAGGGAGATCCCCCACACTGGTACTGAGACACGGACCAGACTCCTACGGGAGGCAGCAGTGAGGAATATTGGTCAATGGGCGCAAGCCTGAACCAGCCATGCCGCGTGCAGGATGACGGTCCTATGGATTGTAAACTGCTTTTGTACAGGAAGAAACCCTCCCTTGTAAGGGAGCTTGACGGTACTGTAAGAATAAGGATCGGCTAACTCCGTGCCAGCAGCCGCGGTAATACGGAGGATCCAAGCGTTATCCGGAATCATTGGGTTTAAAGGGTCCGTAGGCGGTTTTATAAGTCAGTGGTGAAATCTGGTCGCTCAACGATCAAACGGCCATTGATACTGTAAGACTTGAATTACTTGGAAGTAACTAGAATATGTAGTGTAGCGGTGAAATGCTTAGAGATTACATGGAATACCGATTGCGAAGGCAGGTTACTACGAGTATATTGACGCTGATGGACGAAAGCGTGGGGAGCGAACAGGATTAGATACCCTGGTAGTCCACGCCGTAAACGATGGATACTAGCTGTTTGGAGCAATCTGAGTGGCTAAGCGAAAGTGATAAGTATCCCACCTGGGGAGTACGCTCGCAAGAGTGAAACTCAAAGGAATTGACGGGGGCCCGCACAAGCGGAGGAGCATGTGGTTTAATTCGATGATACGCGAGGAACCTTACCAAGGCTTAAATGGGAAACGACAGATTTGGAAACAGATCTTTCTTCGGACGTTTTTCAAGGTGCTGCATGGTTGTCGTCAGCTCGTGCCGTGAGGTGTCAGGTTAAGTCCTATAACGAGCGCAACCCCTGTTGCTAGTTGCCAGCGAGTCATGTCGGGAACTCTAGCAAGACTGCCGGTGCAAACCGCGAGGAAGGTGGGGATGACGTCAAATCATCACGGCCCTTACGCCTTGGGCTACACACGTGCTACAATGGACGGTACAGAGAGCAGCCACTACGCAAGTAGGAGCGAATCTACAAAACCGTTCTCAGTTCGGATCGGAGTCTGCAACTCGACTCCGTGAAGCTGGATTCGCTAGTAATCGCAGATCAGCCATGCTGCGGTGAATACGTTCCCGGGCCTTGTACACACCGCCCGTCAAGCCATGGAAGCTGGGGGTACCTGAAGTCGGTGACCGCAAGGAGCTGCCTTAGGTAAAACTG-3’

Supp. Fig. 2: Partial 16S rRNA sequence of *F. columnare* Fc7. The sequence was obtained by extracting DNA from a pure culture of F. columnare, amplification of the 16S rRNA gene using the primers Eub8F (5’-AGAGTTTGATCMTGGCTCAG-3’) and 1492R (5’-GGTTACCTTGTTACGACTT-3’) and sequencing of the amplicons using Sanger sequencing provided by Eurofins Genomics.

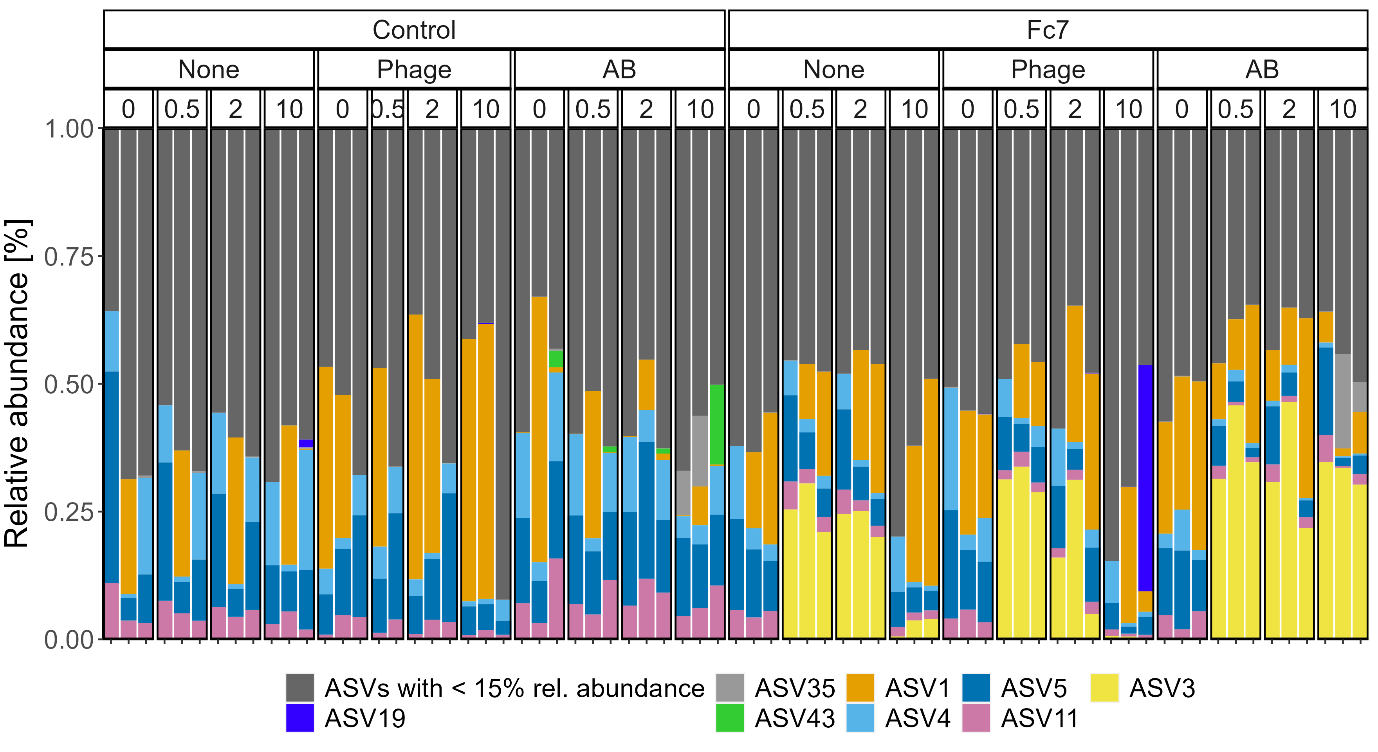

Supp. Fig. 3: Stacked bar graph at ASV level of the bacterial community composition of each water sample taken in Exp.2. All ASVs that do not make up more than 15% of the whole community in at least one sample are summarized as “ASVs with < 15% rel. abundance”. The lowest identified taxonomic group for the ASVs shown are: ASV1: *Polaromonas* sp., ASV3: *Flavobacterium* *columnare* Fc7, ASV4: Comamonadaceae, ASV5: *Janthinobacterium* sp., ASV11: *Janthinobacterium* sp., ASV19: *Pseudomonas* sp., ASV35: Oxalobacteraceae, ASV43: *Flavobacterium* sp.

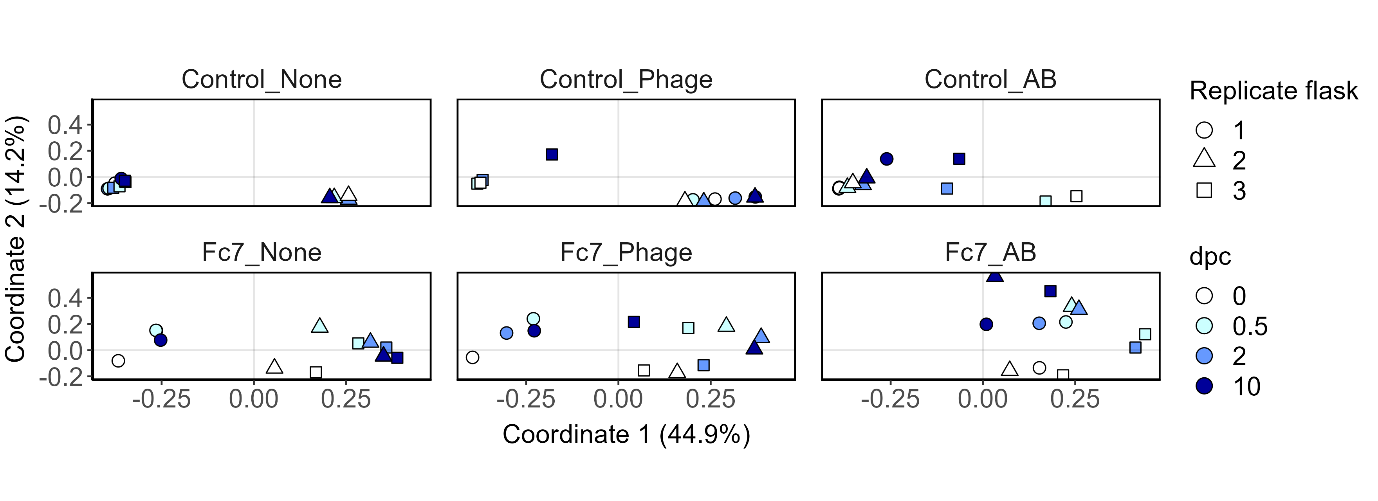
Supp. Fig. 4: PCoA of the Bray-Curtis dissimilarities of all samples from Exp.2. Sampling timepoints are represented by different colours, whereas biological replicate flasks are indicated by different shapes. ASV3 (corresponding to *F. columnare* Fc7) was removed from the dataset prior to generation of the ordination. All panels are from the same PCoA, but were facetted into the different treatment groups.

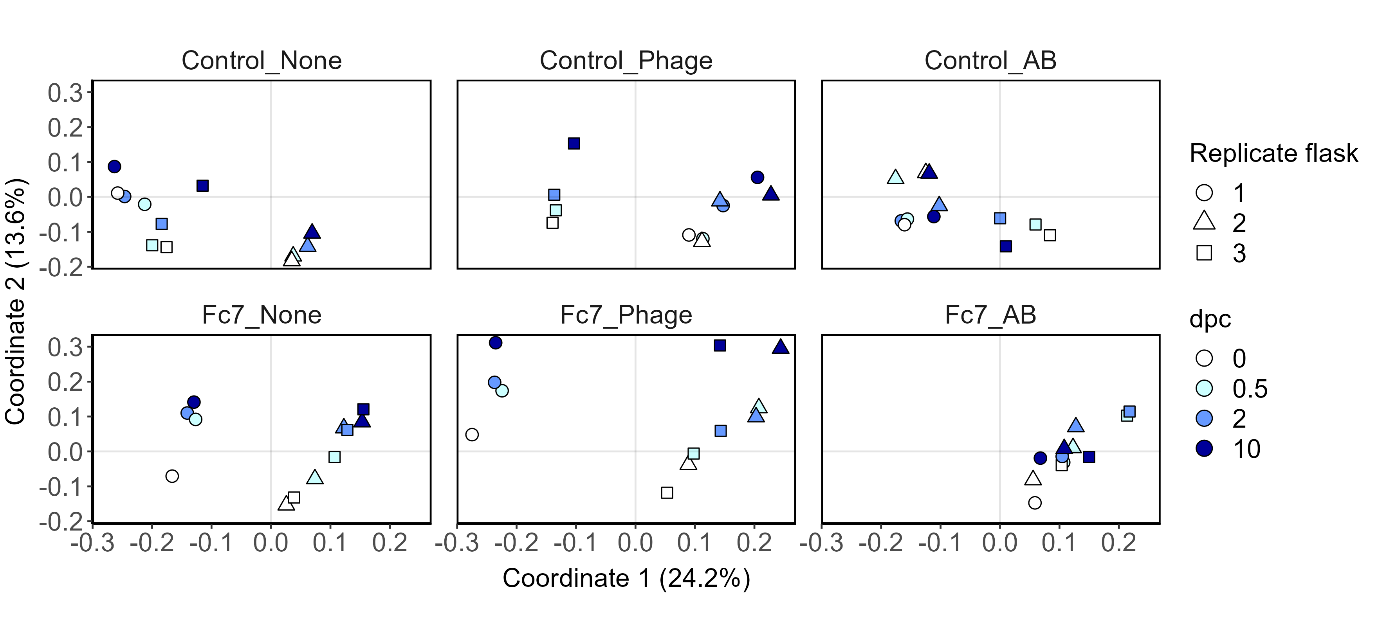

Supp. Fig. 5: PCoA of the Sørensen-Dice indices of the microbiota samples taken in Exp.2. Sampling timepoints are represented by different colours, whereas replicate flasks are indicated by different shapes. Panels are from one PCoA and were facetted into the different treatment groups.

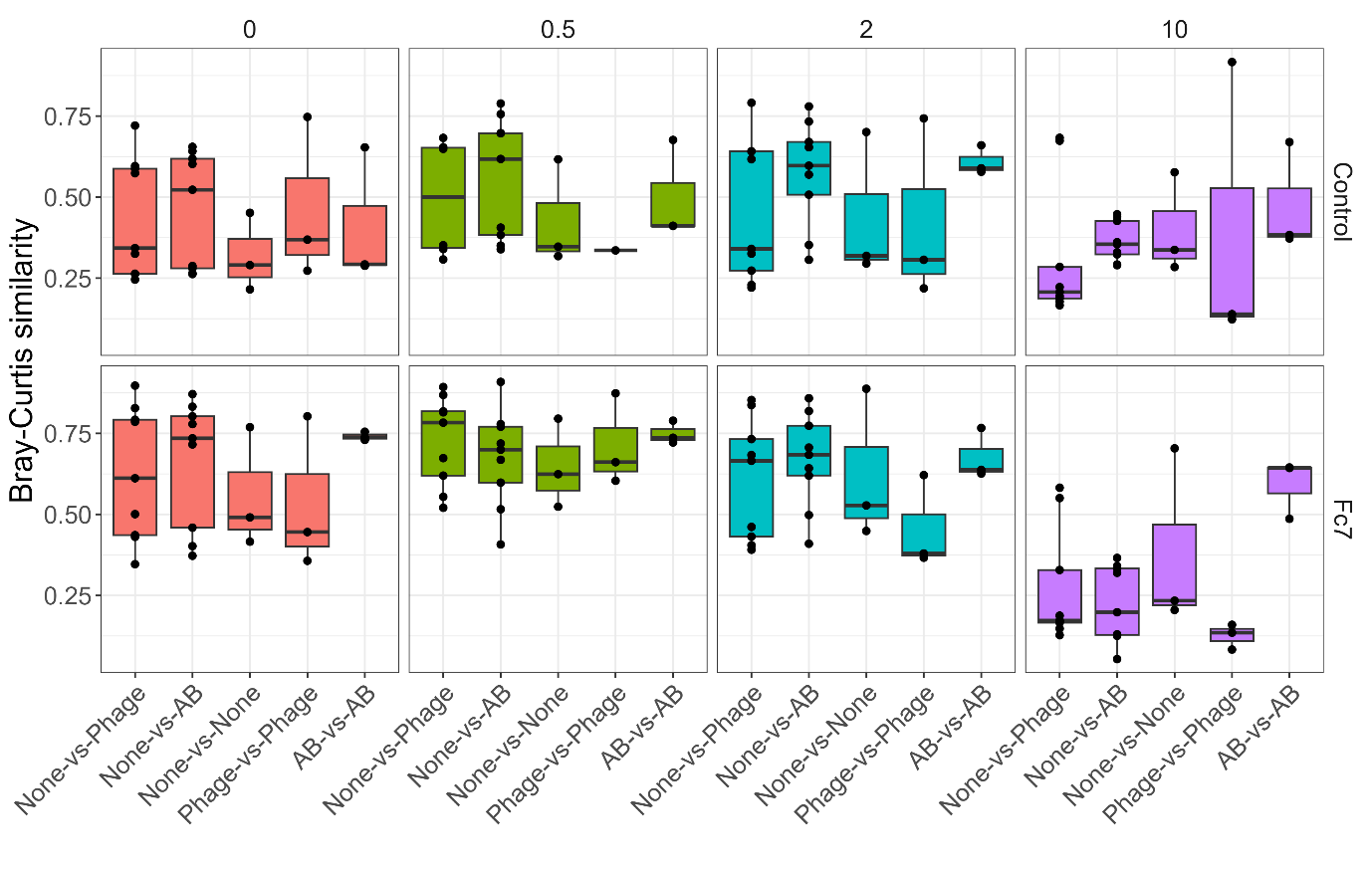

Supp. Fig. 6: Bray-Curtis similarities comparing samples both between and within experimental groups for each timepoint.

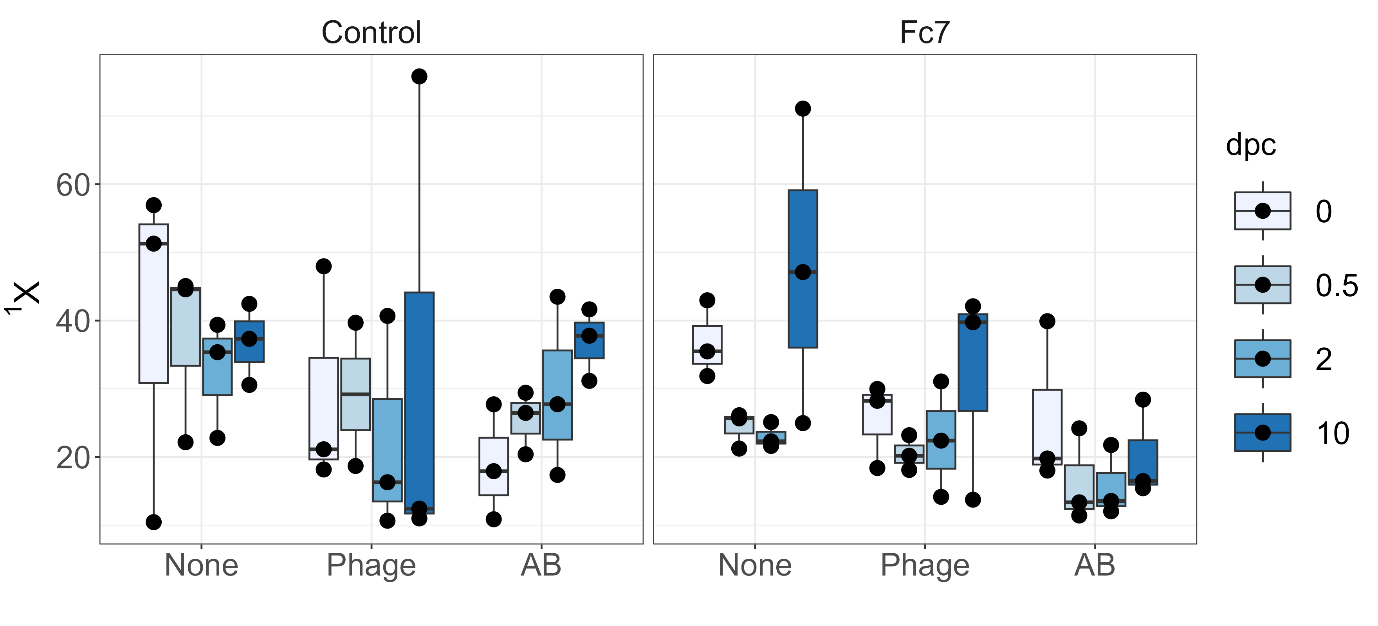

Supp. Fig. 7 α-diversity of all water samples from Exp.2, expressed as Hill’s diversity of order 1 (equivalent to the exponential Shannon index). Box plots show median and upper and lower quartile. Whiskers include all samples.

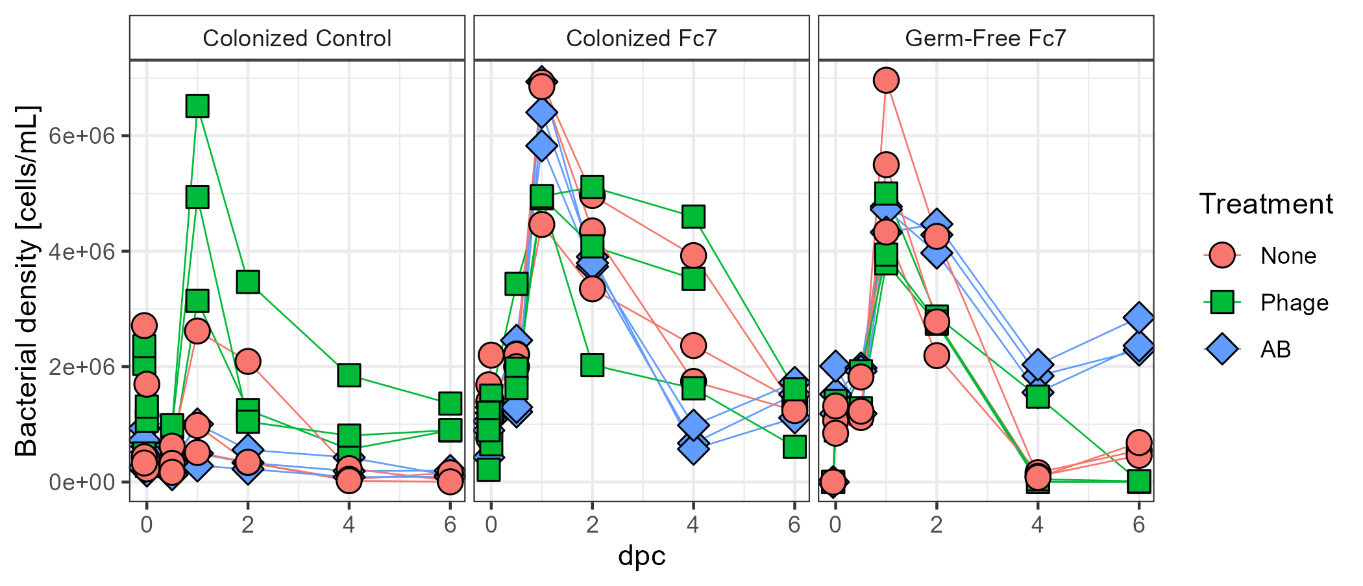
Supp. Fig. 8: The bacterial density [cells per ml] in the rearing water from each flask over time. The samples were taken from the 10 °C experiment and analysed using flow cytometry. At 0 dpc, one sample was taken immediately before and immediately after challenging the flasks with *F. columnare*. At day 1, 4 and 6 the water was exchanged after sampling. Samples from 8 and 10 dpc are not shown due to sample degradation prior to data acquisition. Colours and shapes indicate the treatment type applied at 0 dpc, and the lines connect the bacterial density observations between individual flasks.
